# Systematic Evaluation of AlphaFold2 and OpenFold3 on Protein–Peptide Complexes

**DOI:** 10.64898/2026.04.21.720042

**Authors:** Rumeysa Fayetorbay, Ahmet Can Timucin, Emel Timucin

## Abstract

Protein–peptide interactions are important mediators of diverse biological processes. While deep learning has revolutionized protein structure prediction, comparative evaluation of these methods, specifically for protein–peptide complexes, remains an area of active investigation. Here, we present a systematic benchmarking of AlphaFold2 (AF2) and OpenFold3 (OF3) on a curated, non-redundant dataset of 271 protein–peptide complexes evaluated under CAPRI peptide criteria, partitioned into disordered (IDR) and structured (Non-IDR) peptide subsets. Results show that AF2 consistently outperformed OF3 across both subsets in overall success rate and proportion of high-quality models, while both methods exhibited comparable global fold prediction accuracy. We further demonstrate that AF2 exhibited memorization on a large set of protein–peptide complexes that were in its training data. Analysis of built-in and post-hoc confidence scores demonstrated that PAE-derived metrics, particularly pDockQ2, LIS, and ipSAE, provided the most reliable proxies for structural accuracy in AF2 predictions, whereas OF3’s PAE distributions substantially diminished the discriminative power of its derived scores. Furthermore, we find that canonical DockQ threshold cutoffs for protein–protein complexes are not directly transferable to protein–peptide complexes, underscoring the need for method- and dataset-specific calibration. Peptide sequence composition and length were identified as potential modulators of prediction success, with glycine-rich short peptides and long receptors posing challenges to both methods. Collectively, these findings establish a peptide-specific evaluation framework and highlight the need for dataset/method-calibrated metrics to support the continued development of structure prediction tools for protein–peptide interactions.

## Introduction

Protein-peptide interactions are fundamental to a wide range of biological processes, including cellular signaling events and immune responses. It is estimated that these interactions constitute a large fraction (∼40%) of all macromolecular interfaces within the cell [1–3]. Given their ability to competitively inhibit aberrant protein–protein interactions or mimic peptide hormones, peptides have emerged as important therapeutic agents. This can be exemplified by the clinical success of GLP-1 receptor agonists, which mimic the effects of the endogenous peptide hormone to regulate blood glucose and have transformed the management of type 2 diabetes and obesity [4]. Currently, over 200 peptide-based drugs have been approved by the FDA or are undergoing clinical evaluation [5, 6]. Despite their underscored potential, modeling protein-peptide interactions requires specialized attention due to the enhanced conformational flexibility of peptide ligands [1].

Recent advances in deep learning applications in structural biology, particularly AlphaFold2 (AF2) and the diffusion-based AlphaFold3 (AF3), have transformed the domain by introducing highly accurate predictions [7–9]. Open-source implementations such as OpenFold3 (OF3), Boltz-1, Chai-1, and Protenix have expanded access to these architectures [10–12]. Although these methods have been tested against distinct protein-protein complexes [13–19], their comparative performance for protein-peptide complexes remains relatively underexplored, partly owing to the distinct structural and biophysical properties of peptide partners. These distinctions are epitomized by Short Linear Motifs (SLiMs), which are conserved segments typically 3-15 amino acids in length that govern cellular interaction networks. SLiMs are primarily located within intrinsically disordered regions (IDRs) of proteins [20]. Because the predictive success of methods could be tied to the structured state of the interface, it is essential to evaluate whether the success of deep learning prediction methods is dependent on the structural state of the peptide.

Accuracy of protein-protein complex predictions can be assessed using different criteria. CAPRI classification system assigns models to discrete quality categories (incorrect, acceptable, medium, and high) based on fixed cutoffs for three indicators: *f*_*nat*_, lRMSD, and iRMSD [21]. Alternatively, DockQ provides a continuous score derived from the same three indicators [22, 23]. Because peptides are short, often intrinsically disordered, the structural accuracy criteria established for protein-protein interactions may not adequately reflect accuracy of protein-peptide interactions [24]. In line with this, the CAPRI classification uses a distinct criteria for protein-peptide complexes, where it employs tighter RMSD cutoffs than the standard CAPRI protein–protein criteria to account for the smaller size of peptides [24]. Although well-established DockQ thresholds exist for mapping protein–protein complexes to CAPRI categories [23], the optimal thresholds for protein–peptide complexes remain underexplored, particularly for AF2 and AF3-based methods. In one relevant effort, Xu et al. optimized DockQ thresholds for protein–peptide complexes using MDockPeP2, a global docking program, and reported thresholds that differed substantially from those established for protein–protein complexes [25]. Previous evaluations of deep learning-based protein-peptide structure prediction have primarily focused on AF2 [26–28], with more recent studies expanding to AF3 and other AF3-based methods across datasets of varying sizes [12, 17, 29, 30]. However, none of these studies benchmarking AF2 or AF3/AF3-based methods on protein-peptide complex predictions have utilized a specialized peptide classification system to differentiate structural quality; instead they mostly relied on generic protein-protein DockQ and RMSD metrics. This underscores the necessity of evaluating whether these canonical thresholds remain valid for peptides or if specialized quality standards are required.

In this study, we provide a systematic benchmarking of AF2 and OF3 across three distinct datasets: training complexes (those included in the AF2 training set), as well as IDR (SLiM-like) and Non-IDR (structured peptides). Our findings demonstrate that canonical DockQ thresholds are overly lenient and fail to accurately translate to peptide complexes. Consequently, we propose the adoption of dataset-specific DockQ thresholds to more rigorously evaluate the structural accuracy of protein-peptide interactions. By investigating the memorization effect in AF2 predictions, we demonstrated that performance degrades on structurally novel targets, with the most pronounced decline observed in IDR targets. We further provided an extensive evaluation of PAE-derived metrics for each method and dataset, revealing that discriminative power of these scores was compromised in OF3 due to its narrow PDE distributions. Collectively, our findings established a peptide-specific evaluation to support the rigorous development of these tools.

## Methods

### Dataset Collection

To prepare a comprehensive dataset of experimental protein-peptide complex structures, we used advanced search of RCSB PDB and filtered all hetero-2-mer complexes as of December 2025 and filtered the entries whose one subunit has length ranging between 3 to 15 (inclusive). Given the AF2’s training cutoff date, we split the collected 1230 complexes to “before_Sep2021” (348 entries) or “after_Sep2021” (882 entries) based on the deposition dates of PDB entries. Of these, 882 complexes deposited before the training cutoff of AF2 were designated as training data. To ensure that the remaining complexes were sufficiently distinct from the training set, we clustered the peptide sequences of the entire dataset using MMseqs2 with thresholds of 70% and 50% for identity and coverage, respectively [31]. Within each cluster, the entry with the longest peptide sequence was selected as the cluster representative. We further filtered out any post-cutoff complexes that shared high sequence similarity with training-set entries. This procedure reduced the initial 348 post-cutoff complexes to 282, which were designated as the test set (Fig. 1a, Fig. S1), effectively eliminating sequence and structure redundancy in the final set and providing a more rigorous assessment of model generalization to novel peptide interfaces.

**Fig. 1.**
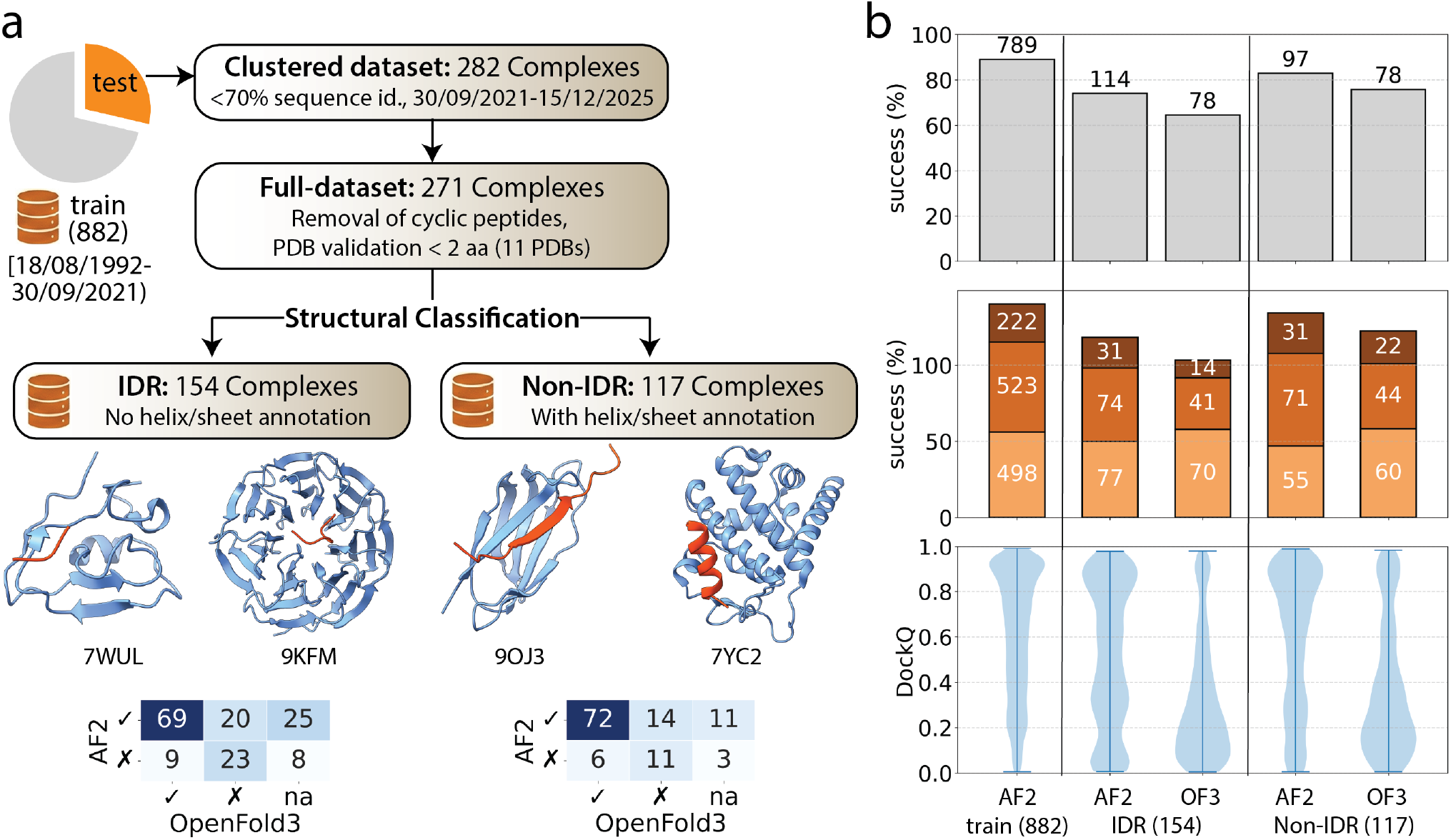
(a) Dataset collection steps and distribution of complexes based on their before/after status according to AF2 training cutoff and clustering peptide sequences with a 70% identity threshold. Confusion matrices show the method’s success based on CAPRI peptide classes. (b) Prediction performance of AF2 and OF3 across datasets. Top plots show the number of correctly predicted complexes based on the CAPRI peptide system. Middle plots show the breakdown of the successful predictions by quality categories (acceptable-light orange, medium-orange, and high-dark orange). Bottom plots show the DockQ score distributions.

We further refined this set by removing cyclic peptides and structures with poor PDB validation, i.e., complexes in which the peptide partner had fewer than 2 resolved amino acids, yielding a final test set of 271 protein–peptide complexes with peptide lengths of 3–15 residues. This set was then split into two subsets based on DSSP secondary structure annotations of the peptide partner. Two example structures from each IDR and Non-IDR set were depicted in Fig. 1a.

### Protein-Peptide Complex Predictions

AF2 v2.3.2 multimer was implemented through ColabFold [8, 32, 33] using MMseqs2 [31] and the UniRef+Environmental database [34]. OpenFold3-preview (v0.3.1) (OF3) [35], an open-source implementation of AF3-based models, was used for structure prediction. For each target, predictions were generated using 5 seeds with 5 samples per seed, yielding 25 models in total.

### Evaluation of Model Accuracy

Accuracy of predicted protein-peptide complexes was assessed using global and interface-based metrics. For global structural accuracy, we have calculated TM-, GDT-TS, GDT-HA and MaxSub scores [36, 37]. For interface metrics, we have used the Critical Assessment of Prediction of Interactions (CAPRI) classification system for protein-peptide criteria [21, 38] and DockQ [22, 23]. CAPRI peptide classification system assigns structural models to discrete quality categories based on specific threshold combinations of three structural similarity metrics: fraction of native contacts (*f*_nat_), ligand RMSD (LRMSD), and interface RMSD (iRMSD). High-quality predictions are defined by *f*_nat_ ≥ 0.8 and either LRMSD ≤ 1.0 Å or iRMSD ≤ 0.5 Å. Medium-quality predictions include either (i) 0.5 ≤ *f*_nat_ *<* 0.8 with LRMSD ≤ 2.0 Å or iRMSD ≤ 1.0 Å, or (ii) *f*_nat_ ≥ 0.8 but with both LRMSD *>* 1.0 Å and iRMSD *>* 0.5 Å. Acceptable models satisfy either (i) 0.2 ≤ *f*_nat_ *<*0.5 with LRMSD ≤ 4.0 Å or iRMSD ≤ 2.0 Å, or (ii) *f*_nat_ ≥ 0.5 with both LRMSD *>* 2.0 Å and iRMSD *>* 1.0 Å. Incorrect models fail to meet any of the above criteria. While the CAPRI system assigns these discrete categories, DockQ score integrates *f*_nat_, LRMSD, and iRMSD into a single continuous score. To determine DockQ thresholds that optimally capture CAPRI peptide quality categories, an exhaustive grid search was performed independently for AF2 and OF3, yielding method-specific threshold sets for each dataset.

### Calculation of Confidence Scores

To assess the confidence of residues located at the interface, we computed interface pLDDT (ipLDDT) scores. For each chain in the structure, residues were identified as interface residues if the distance between their C*α* atoms and the C*α* atoms of residues from any neighboring chain was less than or equal to a defined threshold of 8 Å. Only C*α*–C*α* distances were considered to ensure consistency across all residue types. For each such contacting residue, the corresponding pLDDT score was extracted. Predicted aligned error (iPAE) for AF2 and distance error (PDE) scores were calculated similarly for the interface region. The corresponding interface region in the global PAE/PDE matrix was extracted for each interacting chain pair. PAE values corresponding to these contacting residue pairs were collected, and a normalized interface PAE/PDE score was computed according to the ref. [39]. We also utilized the `ipSAE.py` script to calculate ipSAE, pDockQ, pDockQ2, and LIS [40]. Specifically, ipSAE scores were extracted based on the alignment on peptide. This direction was consistently yielded the maximum ipSAE value across two possible alignment directions, in accordance with the ref. [40].

### Calculation of Physicochemical Properties of Complexes

Several structural and physicochemical properties of the protein–peptide complexes were characterized. Buried surface area (BSA) was calculated as the difference between the sum of solvent accessible surface area (SASA) values of the isolated chains and the SASA of the complex using FreeSASA [41]. Peptide physicochemical properties were computed using `peptides.py` and included hydrophobicity, isoelectric point (pI), aliphatic index, instability index, and Boman index. Charge patterns within the peptide sequences were also assessed using the CIDER tool [42–44], from which we extracted *κ*, along with the fractions of positively and negatively charged residues. Intermolecular hydrogen bonds (H-bonds) between heavy atoms, considering both backbone and side-chain donors and acceptors, and salt bridges between oppositely charged residues were enumerated using custom scripts, applying distance cut-offs of 3.5 Å and 4.0 Å, respectively. Binding free energy (Δ*G*) was estimated using the *PSSM* command of FoldX [45].

## Results and Discussion

### Performance Evaluation Based on Method and Dataset

An initial pool of 1,230 protein–peptide complexes was derived from PDB structures deposited between 1992 and 2025, with peptide lengths ranging from 3 to 15 residues. This dataset was partitioned into three non-redundant subsets: (i) 882 complexes termed training, which fall within the AF2 training data; (ii) 154 complexes termed IDR (SLiM-like), in which the peptide partner lacked any secondary structure; and (iii) 117 complexes termed Non-IDR, in which the peptide contained helical and/or sheet structures (Fig. 1a).

The training set was predicted using AF2, while both the IDR and Non-IDR test sets were predicted using AF2 and OF3. To examine the method agreement, we compared the prediction success of AF2 and OF3 on the IDR and Non-IDR test sets (Fig. 1a-confusion matrices). A complex was considered successfully predicted (✓) if at least one model achieved CAPRI peptide acceptable quality or better; otherwise, it was marked as a failure (×). Targets labeled *na* corresponded to complexes that could not be processed by OF3, limiting its evaluation to 121 of 154 IDR and 103 of 117 Non-IDR complexes. For the IDR set, 69 complexes were successfully predicted by both methods, while 23 were failed by both. Notably, 20 complexes were predicted successfully by AF2 but failed by OF3, compared to only 9 where OF3 succeeded, and AF2 failed, implying an overall performance advantage of AF2. This pattern was also observed for the Non-IDR set, where 72 complexes were jointly successful, 11 failed by both methods, and 14 were uniquely predicted by AF2 against 6 uniquely predicted by OF3. These patterns revealed that although AF2 and OF3 share a broadly overlapping pool of tractable targets, AF2 achieved a larger coverage of the benchmark. Nearly half of the OF3 failures were recovered by AF2, whereas the converse was less common. Furthermore, the consistently higher rate of dual failure in the IDR set relative to the Non-IDR set indicates that intrinsically disordered peptide targets pose an inherently greater challenge to both methods.

We next assessed the success based on CAPRI peptide quality criteria. Top panel in Fig. 1b shows the overall success rate, defined as the percentage of complexes predicted with at least CAPRI peptide acceptable accuracy. AF2 achieved nearly 90% success on the training set (789/882), which is expected given that these structures were included in AF2’s training data. On the test sets, AF2’s success dropped to 74% (114/154) on the IDR set and 83% (97/117) on the Non-IDR set, indicating that structured peptides are more amenable to prediction. AF2 outperformed OF3 on both test subsets, 114 vs. 78 correct targets for IDR and 97 vs. 78 for Non-IDR, reflecting a narrower performance gap between methods for the Non-IDR set. Middle panel in Fig. 1b breaks down prediction performance by CAPRI peptide quality tiers. AF2 produced a larger proportion of high-quality models, particularly on the training and Non-IDR sets than OF3.

We also examined the DockQ distributions and found that AF2 predictions on the training and Non-IDR sets were skewed toward high DockQ values (0.8–1.0), while for both the IDR and Non-IDR test sets, DockQ scores from OF3 models drifted toward lower values (0.0–0.4) (Fig. 1b-bottom). Non-IDR complexes generally yielded higher scores for both methods, aligned with the CAPRI-tier analysis. Overall, these results underscored a consistent performance advantage for AF2 over OF3 in both IDR and Non-IDR datasets for both CAPRI category or DockQ scores. AF2’s overall success (74% IDR, 83% Non-IDR test) aligned with the trend observed by Peng et al., where ∼85% of complexes were predicted with at least acceptable accuracy (DockQ ≥ 0.23) [17]. These success rates were notably higher than those (∼50%) reported by Tsaban et al. [26] and by Ko&Lee [27], both of whom treated the interaction as a folding problem by connecting peptides to receptors via a poly-G linker without MSA information from the peptide. The higher success rate observed on the structured peptide targets in our Non-IDR set was also in line with Tsaban et al.’s finding of lower AF2 success for non-motif-containing peptides compared to motif-containing peptides [26]. Furthermore, our reported success rate of AF2 on its training data was quantitatively consistent with Guan et al.’s observation on 631 complexes, in which they also analyzed AF2’s performance on its training data and showed that AF2 achieved near-90% success [29]. This supported the memorization effect of AF2, suggesting that AF2’s performance degrades for structurally novel targets, which in our study corresponded to the test sets, particularly the challenging IDR subset (Fig. 1b). Moreover, Zhou et al. showed higher performance of AF3-based implementations such as Protenix, Chai-1, and Boltz-1 across 99 complexes compared to AF2 [12]. Contrary to these results, we demonstrated superior performance of AF2 over OF3, which is another AF3-based method, implying a performance gap among AF3-based implementations for protein–peptide complex prediction and in particular an inferiority of OF3-preview (v0.3.1). It is noted that the performance gap observed between AF2 and OF3 may be narrowed by the recent release of OF3-preview2, which showed improved performance relative to the initial version evaluated here.

Despite these consistencies with prior benchmarks, we also observed results that diverged from previous works. Specifically, Johansson-Åkhe and Wallner et al. reported a lower success rate for AF2 (60%, 66 out of 112 complexes) [28] than our corresponding rates of 74% IDR and 83% Non-IDR. Another distinction arose from the comparison with Wang et al. [30], who applied AF2 in a three-stage pipeline and reported a success rate of 34% for top-ranking models. We noted that distinct inference parameters and/or different structural accuracy metrics between this and previous studies could have lead to these discrepancies. Notably, none of the aforementioned studies employed the CAPRI peptide criteria; instead, they mainly relied on DockQ (≥.23) or *f*_*nat*_ (≥.3) or RMSD (≥2 Å) for structural accuracy. Consistent with this point, Zhou et al. reported increased AF2 performance when a more lenient criterion was applied [12], the observation which further signifies the importance of the selected structural accuracy criteria.

In line with AF2’s reduced performance on the IDR subset (Fig. 1b), Tsaban et al. showed that AF2 performed significantly better on peptides with motifs, whereas its performance was notably inferior for non-motif-containing peptides [26]. Furthermore, they demonstrated that AF2 displayed a bias toward predicting alpha-helical structures, noting that helical peptides are modeled particularly well compared to coiled or extended strands. Johansson-Åkhe and Wallner et al. also explicitly analyzed how disorder affects prediction, concluding that highly disordered peptides yielded a lower average DockQ score—albeit not statistically significant—compared to ordered peptides [28]. Additionally, Wang et al. [30] tested AF2 on 62 complexes with fully disordered peptides that form random coils; they showed that a three-stage pipeline, utilizing AF2 for multimer prediction followed by ZDOCK rigid docking and refinement using the AMBER force field and machine learning potentials, performed better at predicting these challenging random-coil peptide partners.

### Optimal DockQ Thresholds Capturing CAPRI Peptide Criteria

While protein-protein quality is typically assessed via CAPRI categories or DockQ scores, these metrics may fail to account for the protein-peptide interfaces, necessitating specialized cutoffs to accurately map predictions to CAPRI quality levels. To address this gap, we determined optimal DockQ thresholds for distinguishing between CAPRI peptide quality categories for both AF2 and OF3 across all datasets using grid optimization (Table 1).

**Table 1.** Optimal DockQ thresholds for CAPRI peptide by method and dataset.

| dataset | method | I\|A | A\|M | M\|H | accuracy | accuracy* |
| --- | --- | --- | --- | --- | --- | --- |
| train | AF2 | 0.448 | 0.720 | 0.924 | 0.828 | 0.795 |
| IDR | AF2 | 0.409 | 0.711 | 0.912 | 0.861 | 0.857 |
|  | OF3 | 0.409 | 0.711 | 0.913 | 0.926 | 0.924 |
| Non-IDR | AF2 | 0.447 | 0.651 | 0.922 | 0.848 | 0.816 |
|  | OF3 | 0.445 | 0.782 | 0.916 | 0.897 | 0.874 |

I: incorrect, A: acceptable, M: medium, H: high
\*Accuracy for DockQ thresholds of 0.4, 0.7, 0.9 from the ref. [25].

Both AF2 and OF3 yielded similar DockQ boundaries for the IDR dataset, converging on approximate thresholds of 0.41 for the incorrect–acceptable boundary, 0.71 for acceptable–medium, and 0.91 for medium–high. Notably, these optimized thresholds closely aligned with the previously proposed thresholds of 0.4, 0.7, and 0.9 for MDockPeP2 [25], and both threshold sets yielded comparable accuracies (Table 1, Fig. S2), supporting the validity of the DockQ thresholds established in MDockPeP2 [25] for the IDR dataset. However, optimal DockQ thresholds for the Non-IDR dataset deviated from those of the IDR dataset as well as from the previously established thresholds [25]. Specifically, the incorrect–acceptable boundary was 0.45 and the medium–high boundary was 0.92 for both AF2 and OF3, while each method produced notably different thresholds for the acceptable–medium boundary, which is 0.65 for AF2 and 0.78 for OF3.

Several aforementioned protein-peptide benchmarking studies exclusively used DockQ with standard protein-protein cutoffs (0.23, 0.49, 0.80) for structural accuracy assessment [12, 28, 29]. Our DockQ threshold optimization against CAPRI peptide criteria revealed a caveat regarding the appropriateness of applying protein-protein DockQ cutoffs directly to protein-peptide complexes (Table 1). Specifically, the optimal DockQ boundaries for distinguishing CAPRI peptide quality categories deviated substantially from canonical protein-protein values for both IDR and Non-IDR sets. This dependence of optimal thresholds on structural content implies that success rates reported under protein-protein cutoffs would lead to inflated results that are difficult to compare across studies. Accordingly, using the generic DockQ threshold of 0.23 would have led a large fraction of CAPRI-incorrect complexes to be classified as correct for both methods and all datasets (Fig. S2), resulting in an apparent inflation of the success rates reported here (Fig. 1b). We further noted that while the thresholds established for the IDR dataset closely aligned with those reported by Xu et al. [25], the Non-IDR set showed different thresholds, further extending the previous studies for protein-peptide complexes with structured peptides. Overall, this analysis underscores the dependence of DockQ thresholds on the structural content of the predicted peptide, and partly on the method, as seen in the OF3 example, highlighting the need to evaluate thresholds on a per-dataset/method basis before adopting them for protein-peptide complex assessment.

Consequently, protein-peptide benchmarks relying solely on protein-protein DockQ thresholds should be interpreted with caution, and dataset-specific threshold calibration might be essential before drawing conclusions about predictive success in protein–peptide complex predictions. Taken together, these findings extended the prior evaluations by deriving a peptide-specific DockQ thresholds, reflecting that the success rates may shift notably when evaluation criteria are appropriately tailored to the protein–peptide targets.

### Comparison of Model Confidence Scores and Interface Properties

We next assessed method agreement based on global and interface-based metrics using the models predicted by both methods. Among interface-level metrics (Fig. S3-top panels), DockQ and *f*_nat_ showed weak agreement between AF2 and OF3 predictions, whereas iRMSD and LRMSD were more strongly correlated. Interface metric agreement was consistently lower in the Non-IDR subset (Fig. S3b) compared to the IDR subset (Fig. S3a). Overall, interface metrics exhibited considerable scatter, with regression lines (Fig. S3-red lines) falling below the diagonal, indicating that AF2 consistently produces higher-quality complex-level predictions (Fig. S3-top rows). This trend was further reflected in the proportion of data points with higher AF2 scores as such 62–73% of complexes achieved higher DockQ and *f*_nat_ values in AF2. The bottom rows of Fig. S3 present global structural similarity metrics. In contrast to interface metrics, these scores were closely aligned between AF2 and OF3 for both subsets (*r >* 0.87), indicating that both methods predict the overall fold with comparable accuracy. Taken together, these analyses revealed strong global fold agreement between methods albeit a weak one for interface, implying that the primary challenge, particularly for OF3, lies in accurate peptide placement, rather than in structure prediction.

We also assessed method agreement based on model confidence scores and physical properties of the complexes from the IDR (Fig. 2a) and Non-IDR subset (Fig. 2b). Among the built-in confidence scores, pTM showed the strongest inter-method correlation in both subsets (*r* = 0.57– 0.75), while ipTM and model confidence exhibited weaker agreement for IDR (*r* = 0.38), with the Non-IDR subset showing even weaker correlations (*r* = 0.17–0.18). Across all confidence metrics except for pLDDT, regression lines (black dashed) were positioned above the diagonal (below the diagonal for iPAE), indicating that OF3 consistently produced higher confidence scores than AF2 for both subsets. This pattern was most pronounced for iPAE, where only 2% of data points showed higher scores in AF2 models in both subsets, suggesting that OF3 extremely overestimates confidence scores, particularly PDE in OF3, relative to AF2. Notably, iPAE scores showed near-uniform AF2 dominance, with 98% of data points exhibiting higher AF2 iPAE values, i.e., lower confidence, in both subsets, corroborating OF3’s tendency toward inflated interface confidence scores despite its lower success than AF2 (Fig. 1).

**Fig. 2.**
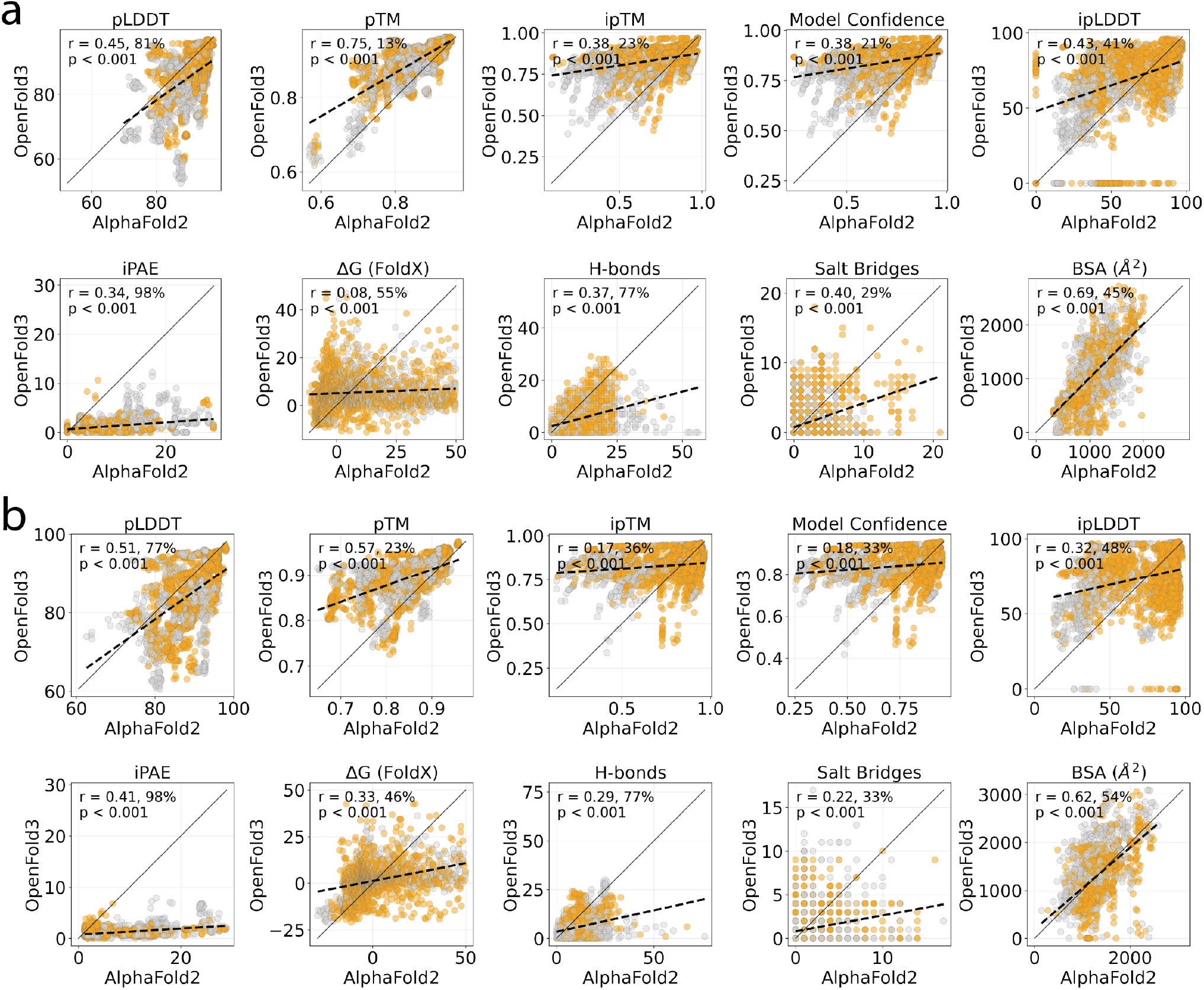
Comparison of confidence scores and complex physical properties from AF2 versus OF3 (a) IDR and (b) Non-IDR subsets. Pearson correlations (r) and proportion of data points with higher AF2 scores than OF3 scores were also given. Gray color indicates incorrect models and orange indicates CAPRI acceptable+.

To examine the relationship between built-in confidence scores and structural accuracy for each method and dataset, we plotted each confidence metric against DockQ across all predictions (Fig. S4). Global confidence scores, pLDDT and pTM, exhibited uniformly weak correlations with DockQ for all subsets including the training set predicted by AF2, confirming their limited utility as proxies for DockQ-based quality. In contrast, confidence scores derived from interface including ipTM, model confidence, and ipLDDT, generally showed stronger correlations. OF3 showed consistently lower correlations across these same scores in both subsets, indicating diminished discriminative power of OF3’s scores relative to AF2. The most pronounced discrepancy was observed for ipLDDT in the IDR subset, where OF3’s correlation dropped to *r* = 0.33 compared to AF’s 0.59, further consolidating that OF3’s interface confidence scores were particularly less reliable for IDR-containing peptide targets. Together, these results established ipLDDT and ipTM as the most informative confidence metrics for DockQ-based filtering, and revealed that confidence scores are systematically less discriminative for OF3, particularly on IDR targets.

We further characterized each predicted interface by computing physical properties including the number of hydrogen bonds (H-bonds), salt bridge interactions, and total buried surface area (BSA), alongside the binding free energy (ΔG) estimated using FoldX [45]. Among these physical features, BSA displayed the strongest inter-method correlation (*r* = 0.62– 0.69) with roughly balanced distributions for both subsets (Fig. 2). AF2 predicted a greater number of interface hydrogen bonds in 77% of cases across both subsets. Salt bridges, by contrast, showed the opposite pattern, with OF3 yielding higher counts in the majority of cases (*r* = 0.40, 29% AF2-higher in IDR). Δ*G* values showed the weakest correlation in the IDR subset (*r* = 0.08), improving in the Non-IDR subset (*r* = 0.33), and exhibited high scatter throughout, reflecting substantial disagreement in predicted binding free energies. Altogether, these results indicated that while both methods assigned comparable global fold confidence (Fig. S4), they diverged considerably in interface-level confidence estimation and physicochemical interface characterization.

We also evaluated how well each method reproduced these physical properties of the interface (BSA, number of H-bonds and salt bridges) relative to the corresponding native complex structures (Fig. S5). Across both subsets, BSA showed the strongest correlation with crystal structure values for both methods. In the IDR subset, AF2 achieved *r* = 0.67, outperforming OF3, with OF3 exhibiting notably larger error magnitudes despite comparable trends (Fig. S5a). In the Non-IDR subset, AF2’s BSA correlation improved to *r* = 0.73, whereas OF3 showed a slight decline (Fig. S5b), further widening the gap between methods on structured targets. H-bond counts were reproduced with low fidelity by both methods, with near-identical correlations in the IDR subset and a small AF2 advantage in the Non-IDR subset. Regression lines for OF3 H-bond predictions fell consistently below the identity diagonal, particularly in the Non-IDR subset, indicating a systematic tendency to underestimate hydrogen bond counts relative to crystal structures. Salt bridge reproduction was the weakest across all conditions and revealed the largest inter-method divergence. In the IDR subset, OF3 showed markedly poorer performance than AF2, with its regression line substantially below the diagonal, reflecting consistent underestimation of salt bridge counts. In the Non-IDR subset, both methods exhibited weaker salt bridge correlations than in the IDR set, with comparable errors (Fig. S5b). Taken together, these results indicate that AF2 more reliably recapitulated the physical properties of the interface across both disordered and structured targets, with OF3 showing a consistent pattern of underestimating interface contacts. Collectively, the reproduction of interface physical properties from crystal structures was weak at best across all conditions, with BSA consistently yielding the strongest correlations. Notably, both methods performed better on Non-IDR targets for BSA, whereas H-bond and salt bridge reproduction did not improve substantially or even slightly deteriorated on structured targets relative to IDR ones, indicating that the inherent flexibility of disordered peptides may not the sole source of physical property prediction error of AF2 or OF3. In line with this notion, Johansson-Åkhe and Wallner evaluated electrostatic and shape complementarity and found that AF2 tends to lack high electrostatic complementarity at protein-peptide interfaces, proposing that the raw AF2 outputs may lack proper physical refinement [28].

### Comparison of Post-hoc Confidence Scores

AF2’s native ipTM score, while recognized as a proxy for prediction quality [13], is known to be sensitive to the length and disorder content of input sequences [40]. When full-length sequences containing disordered regions or accessory domains are provided, ipTM is diluted by residue pairs that do not contribute to binding, leading to underestimation of true interface confidence. This limitation could be particularly critical for protein-peptide complexes where the peptide contains disordered regions [46, 47]. To address this, Bryant et al. introduced pDockQ, based on interface pLDDT and contact counts [18], which was extended to pDockQ2 by incorporating PAE values for interface residue pairs [39]. Kim et al. further proposed the Local Interaction Score (LIS), which averages PAE-derived pTM values over interchain pairs within 12 Å[48]. More recently, Dunbrack introduced ipSAE, which addresses the length-dependency of ipTM by filtering interchain residue pairs by a PAE cutoff and simultaneously rescaling the *d*_0_ parameter of the TM-score equation to the number of residues passing the filter, yielding scores that are robust to disordered flanking sequences [40]. Varga et al. also derived another ipTM-based scores, actifpTM, which uses predicted distance distograms to restrict ipTM to predicted interface [49]. To assess how well these post-hoc confidence scores reflect structural accuracy, we also evaluated their correlations with DockQ (Fig. 3).

**Fig. 3.**
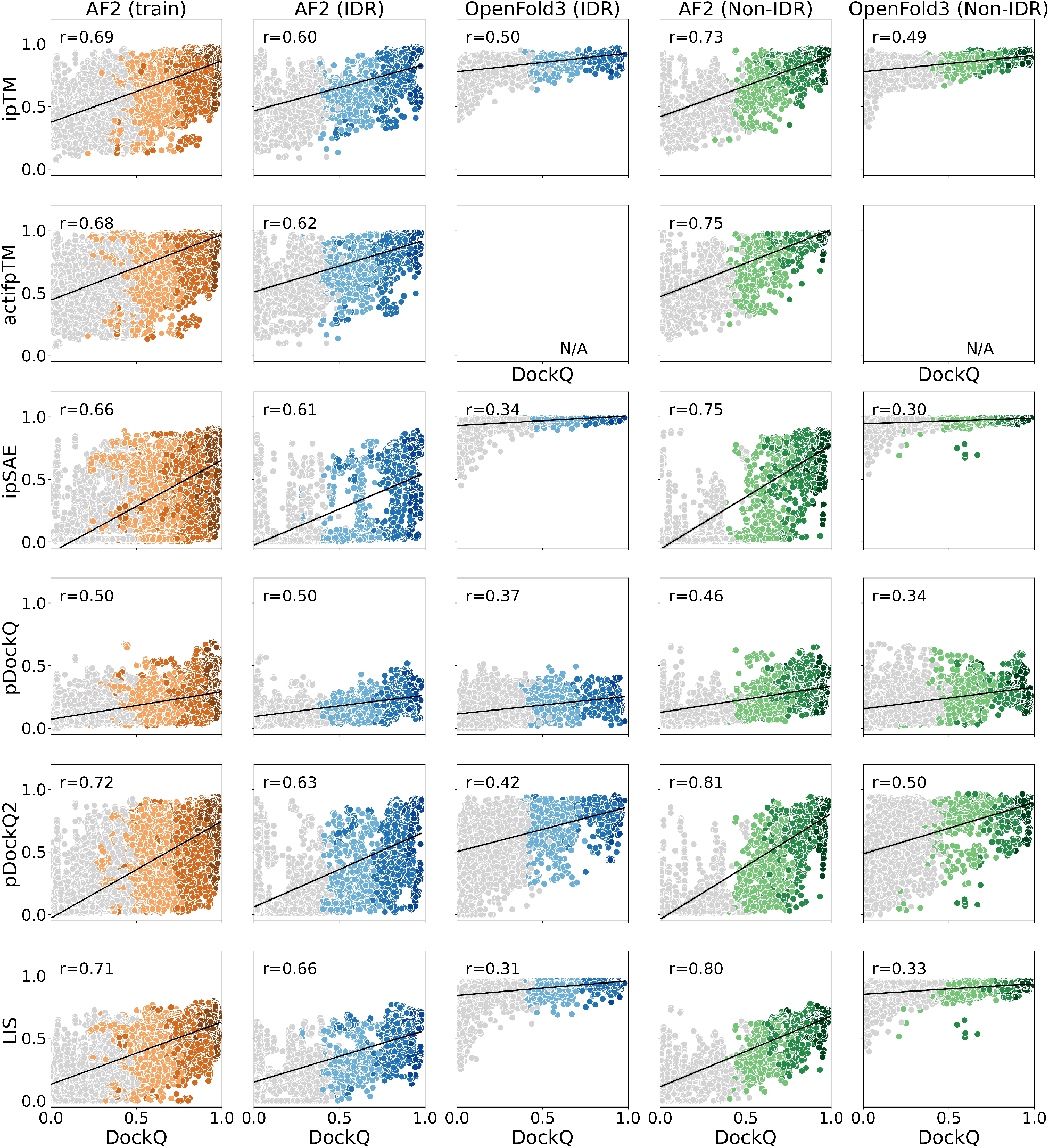
Distribution of post-hoc confidence scores across DockQ. Dots were colored according to CAPRI peptide classes from acceptable (light) through medium to high (dark). Incorrect models are shown in gray.

Correlations were consistently higher for AF2 than for OF3 across both IDR and Non-IDR subsets, and higher for Non-IDR than IDR within AF2, indicating that structured peptide interfaces are more amenable to scoring by these scores. This pattern was particularly pronounced for OF3 on the IDR subset, where all metrics showed markedly lower correlations with DockQ relative to their AF2 counterparts (Fig. 3). pDockQ yielded the weakest correlations across all conditions, suggesting that interface pLDDT alone, without PAE information, is insufficient to capture structural accuracy in protein-peptide complexes. Across AF2 predictions, pDockQ2 and LIS consistently ranked as the strongest correlates of DockQ, with ipSAE and actifpTM showing comparable performance to one another. Notably, the gains of these specialized metrics, except for pDockQ, over ipTM in AF2 predictions was the most apparent in the test sets as they slightly improved correlations. On the other hand, a ceiling effect was evident in several OF3 panels, where these scores clustered near their maximum values regardless of DockQ distribution. This behavior is consistent with the compressed iPDE distributions observed for OF3 for both subsets, in which predictions were assigned high confidence (Fig. 2). Thus, PAE-derived scores lost their discriminative power for OF3, meaning that the rationale behind developing these metrics does not transfer to OF3 predictions. Taken together, these results indicated that while pDockQ2, LIS, and ipSAE provide the most reliable confidence proxies for AF2-based protein-peptide predictions, the utility of all evaluated PAE-derived scores was substantially lower for OF3 predictions, implying that ipTM remains the most practical metric for OF3 predictions.

To evaluate the ability of each score to distinguish correctly predicted complexes from incorrect predictions (CAPRI peptide acceptable+ vs incorrect), we computed AUROC and normalized AUPRC for all metrics across all datasets (Fig. 4). Normalized AUPRC accounted for the different positive rates across subsets, mapping random performance to zero and perfect classification to one, enabling direct comparison across different subsets. Among all, ipSAE and LIS ranked as the top-performing scores for AF2 predictions of IDR subset, with both scores outperforming the baseline ipTM and pTM scores in both AUROC and normalized AUPRC. These two stayed among the top performers, whilst pDockQ2 and actifpTM joined them as showing the highest AUROC and normalized AUPRC in the Non-IDR subset. Although we reported a comparable level of DockQ correlations of the post-hoc scores with raw ipTM score (Fig. 3), these results confirm that restricting scoring to interface-proximal residues, as done by ipSAE, pDockQ2, actifpTM, and LIS, provided a meaningful advantage over whole-chain ipTM for protein-peptide complex evaluation.

**Fig. 4.**
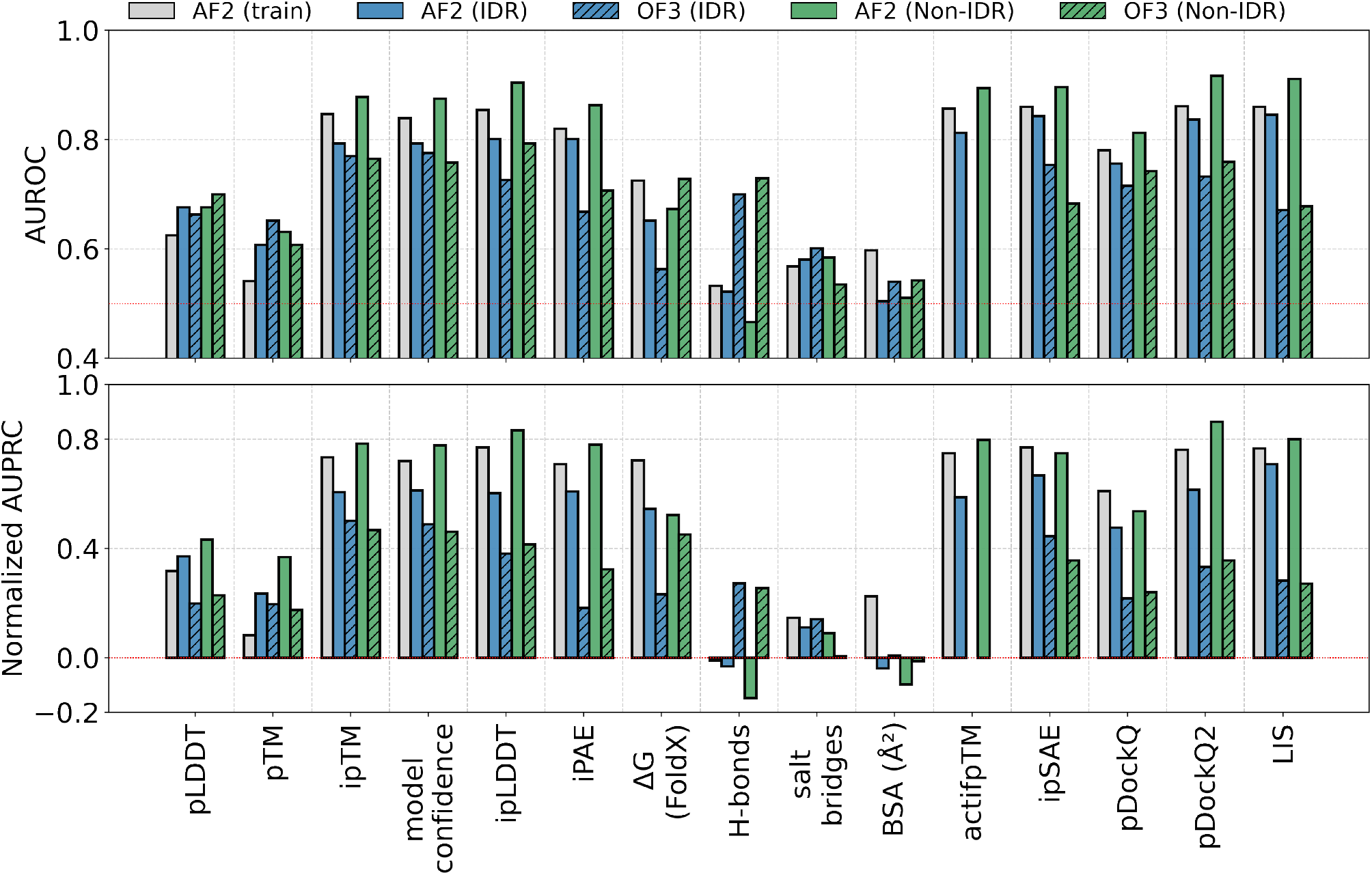
Binary classification performance of tested scores and physical complex features based on AUROC and normalized AUPRC.

Among the physical interface features, ΔG showed weak discriminative ability, ranking below the confidence scores, while H-bonds, salt bridges and BSA showed near-random performance, indicating no predictive value for AF2 and OF3.

### Dissecting the Success and Failure Modes Based on Peptide Features

To investigate whether or not AF2 or OF3 has a tendency towards a given peptide sequence, we determined a number of sequence-level determinants and assessed the prediction success based on these determinants. First, we found that success rates in the training were uniformly high across all peptide lengths (0.85-0.94), with no length showing a pronounced failure bias, consistent with the expectation that these interactions were well-represented in AF2’s parameters (Fig. S6). On the other hand, lower and more variable success rates were reported for the test sets; yet the success rates were not monotonically related to length (Fig. 5-top left).

**Fig. 5.**
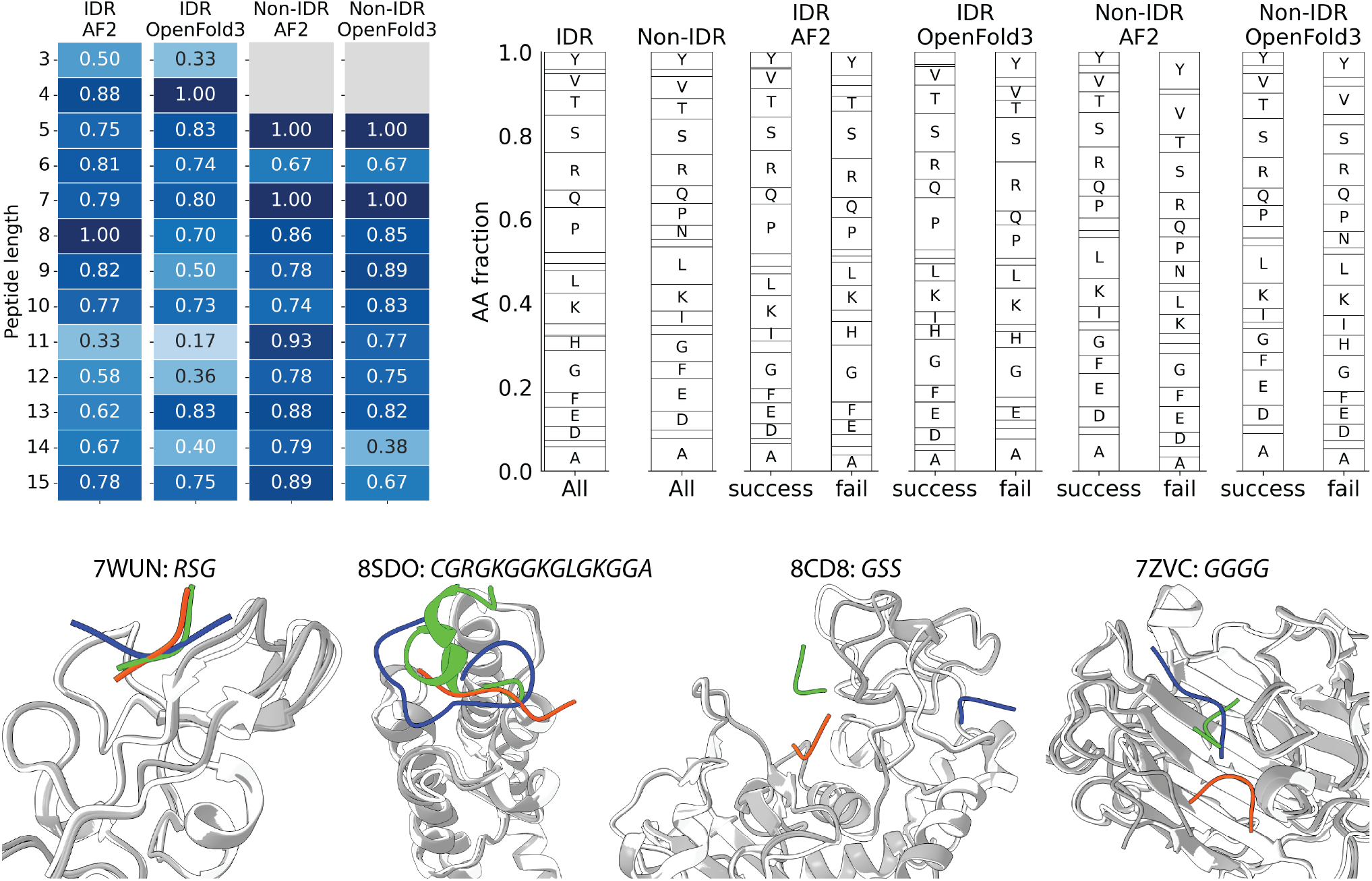
(Left) Success rates at different peptide lengths. (Right) Mean amino acid fractions in the entire dataset (All), successful and failed predictions. Representative crystal structures (orange-white) with AF2 (blue-gray) and OF3 (green-gray) predictions are displayed for four incorrect predictions.

This observation was in line with the previous reports on AF2, that did not find any association between peptide length and prediction success of AF2 or OF3 [26–28].

We also assessed whether or not composition of amino acids have any effect on prediction success by extracting the amino acid fractions of the peptides from each successfully and failed predictions and reported the average fractions in Fig. 5-top right. Composition of training set peptides was broadly similar between successful and failed predictions (Fig. S6). However, failed predictions showed a slightly enriched H containing peptides. Compositions of the IDR and Non-IDR subsets were also broadly similar with subtle differences observed such that the IDR subset contained higher P (≥ 5%), consistent with the known composition of flexible and disordered peptides [50], while the peptides in the Non-IDR subset were enriched in E and L (≥ 5%). Failed predictions in IDR subset tended to be enriched in G for AF2 and R for OF3. For the Non-IDR subset, the failed predictions showed enrichment of V and Y for AF2 and successful predictions showed A enrichment, while OF3 predictions did not show any notable enrichment (≥ 5%) in either successful and failed predictions. These compositional differences, albeit subtle, were not consistent between AF2 and OF3, suggesting that sequence composition-related challenges may be related to the method.

Representative crystal structures and corresponding AF2 and OF3 predictions are shown for four entries (Fig. 5, bottom). All of these visualized cases were incorrect predictions according to the CAPRI peptide system. Among these, the target 8SDO (*CGRGKGGKGLGKGGA*) and 7ZVC (*GGGG*) highlighted the challenges posed by G-rich and repeat-sequence peptides, where both methods failed to reproduce the binding pose, consistent with the compositional analysis showing enrichment of G in failed predictions. 7WUN (*RSG*) and 8CD8 (*GSS*) further illustrated the challenge of very short peptides.

In the AF2 Non-IDR subset, length ratio and receptor length showed statistically significant differences between groups, with failed predictions associated with higher length ratios and shorter receptor lengths (*p <* 0.05). In the OF3 Non-IDR subset, receptor length again showed a significant difference, suggesting that complexes with long receptors and structured peptides posed a greater challenge for both methods. While other features including hydrophobicity and charge-related features were less discriminative in both sets and for both methods. In the training set, physicochemical features were broadly similar between successful and failed predictions, with the most notable, but not significant, difference observed in *κ*, where failed predictions showed wider distributions shifted toward higher values, indicating greater charge asymmetry in failed peptides (Fig. S6). For the IDR subset, the failed predictions for both methods exhibited wider *κ* distributions (Fig. S7). For the Non-IDR subset, failed predictions from AF2 showed larger *κ* values while those from OF3 showed slightly lower *κ* values (Fig. S7). Taken together, length-related features emerged as the most consistent discriminators for structured peptides for both methods, while no other physicochemical features provided a significant separation between success and failure for IDR peptides.

According to previous reports, length, either of the peptide or the receptor, did not have significant impact on the prediction accuracy of AF2, particularly DockQ [26–28]. In fact, this length independence has been highlighted as an advantage of deep learning-based prediction methods over traditional docking tools, where sequence length was known to exert an effect on the success of complex modeling [1, 51]. Despite these reports, our findings showed that particularly for structured peptides, receptor length could be a factor impacting the prediction success of both AF2 and OF3, while for disordered peptides we did not report a statistically significant effect of length on the success. Overall, based on these assessments, we noted that despite the absence of a significant correlation between peptide length and prediction success (Fig. 5), large receptors, especially when co-crystallized with shorter peptides, i.e., lower peptide-to-receptor length ratios, pose a challenge for AF2 and OF3, as for traditional docking tools.

## Conclusions

Rapid release of new structure prediction methods has arguably outpaced systematic evaluation efforts, leaving the relative strengths and limitations of these methods for peptide targets less characterized. Thus, a comprehensive comparative analysis provided here for protein-peptide targets offered the following advantages. First, structural accuracy and confidence metrics embedded in these models were mainly developed and validated for protein-protein interfaces, and their transferability to peptide partners could not be assumed. Second, the structural diversity of peptide binding modes, ranging from fully disordered recognition motifs to *α* and *β*-containing interfaces, demanded that evaluations account for this heterogeneity rather than reporting aggregate statistics that may obscure method-specific failure modes. Third, training data cutoffs of different methods may vary, and the extent to which reported success rates reflect generalization versus memorization of training set structures needs to be separated. Addressing these gaps, our study provides a systematic benchmarking of AF2 and OF3 for protein–peptide complex structure prediction across disordered (IDR), and structured (Non-IDR) peptide containing datasets evaluated under CAPRI peptide criteria. Taken together, these findings underscore the need for peptide-specific evaluation frameworks and method-calibrated confidence metrics for the continued development and rigorous assessment of structure prediction tools for protein–peptide interactions.

## Supporting Information

Figures (S1–S7) detailing dataset construction, comparative structural performance, confidence score distributions, and physicochemical features of prediction success.

## Data and Software Availability

All predicted structures including native PDB structures are available at https://zenodo.org/records/19598102. Analysis scripts are available at https://github.com/timucinlab/SLiM_preds/.

## Author Contributions

E.T. and A.C.T conceived and supervised the study. Data curation was performed by R.F. and A.C.T., Formal analysis was conducted by E.T. and R.F. All authors contributed to the writing and final editing of the manuscript.

## Notes

### Competing Interest Statement

The authors have declared no competing interest.

