## Supporting Information for "Systematic Evaluation of AlphaFold2 and OpenFold3 on Protein–Peptide Complexes"

### Dataset Construction

We compiled a non-redundant dataset of 1,230 SLiM complexes, of which 882 predated September 30, 2021, corresponding to the AF2 training cutoff date, while 348 were released after this date (Fig.S1a). We also performed sequence clustering of SLiM sequences at 70% identity with 50% coverage to ensure that the test set does not contain any similar SLiM sequences to those found in AF2's training dataset (Fig. S1a). Clustering reduced the 1,230 complexes to a set of 987 SLiM complexes, effectively reducing redundancy in the final test set of 282 post-AF2 cutoff complexes (Fig. S1b). The clustered dataset comprised 987 structures, predominantly determined by X-ray crystallography (843), with additional contributions from cryo-EM (140) and NMR spectroscopy (4) (Fig. S1b). Resolution distribution showed a peak around 2 Å, with most structures falling between 1.5 and 2.5 Å (Fig S1b). Receptor proteins exhibited considerable size variation, with lengths ranging from approximately 50 to over 1000 amino acids, showing a right-skewed distribution (Fig. S1b). SLiM lengths displayed a distribution centered around 8-10 amino acids, consistent with the typical length range of linear motifs (Fig. S1b).

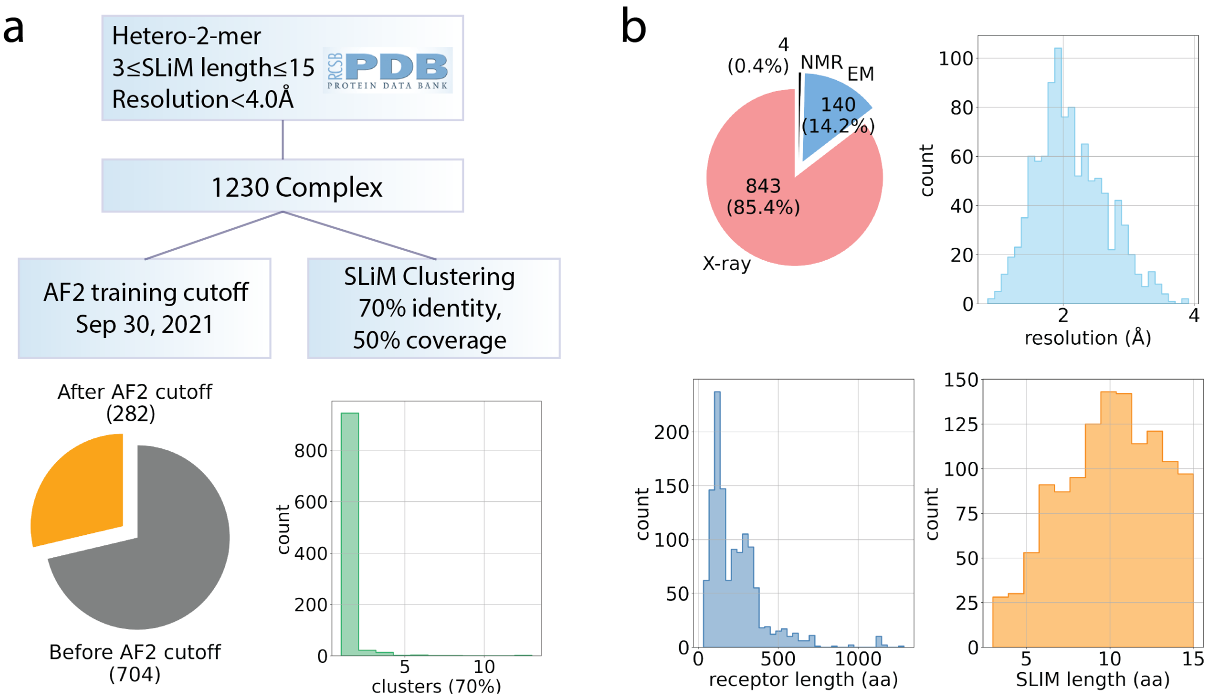


**Fig S1.** (a) Dataset collection steps including clustering of the peptide sequences of all collected complexes using a 70% identity threshold. (b) Distribution of experimental method and resolution of full-dataset. Bottom plots depict receptor and peptide (SLiM) lengths.

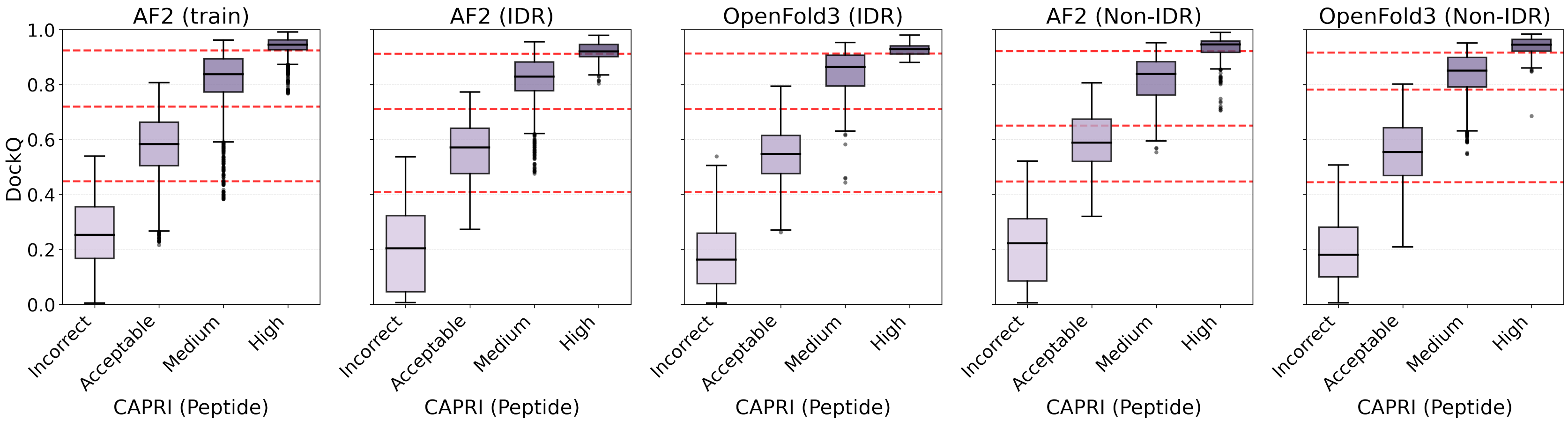


**Fig S2.** Distribution of DockQ scores across CAPRI peptide classes for each method. Red dotted lines indicate the 0.4, 0.7, and 0.9 DockQ thresholds tailored for protein-protein complexes.

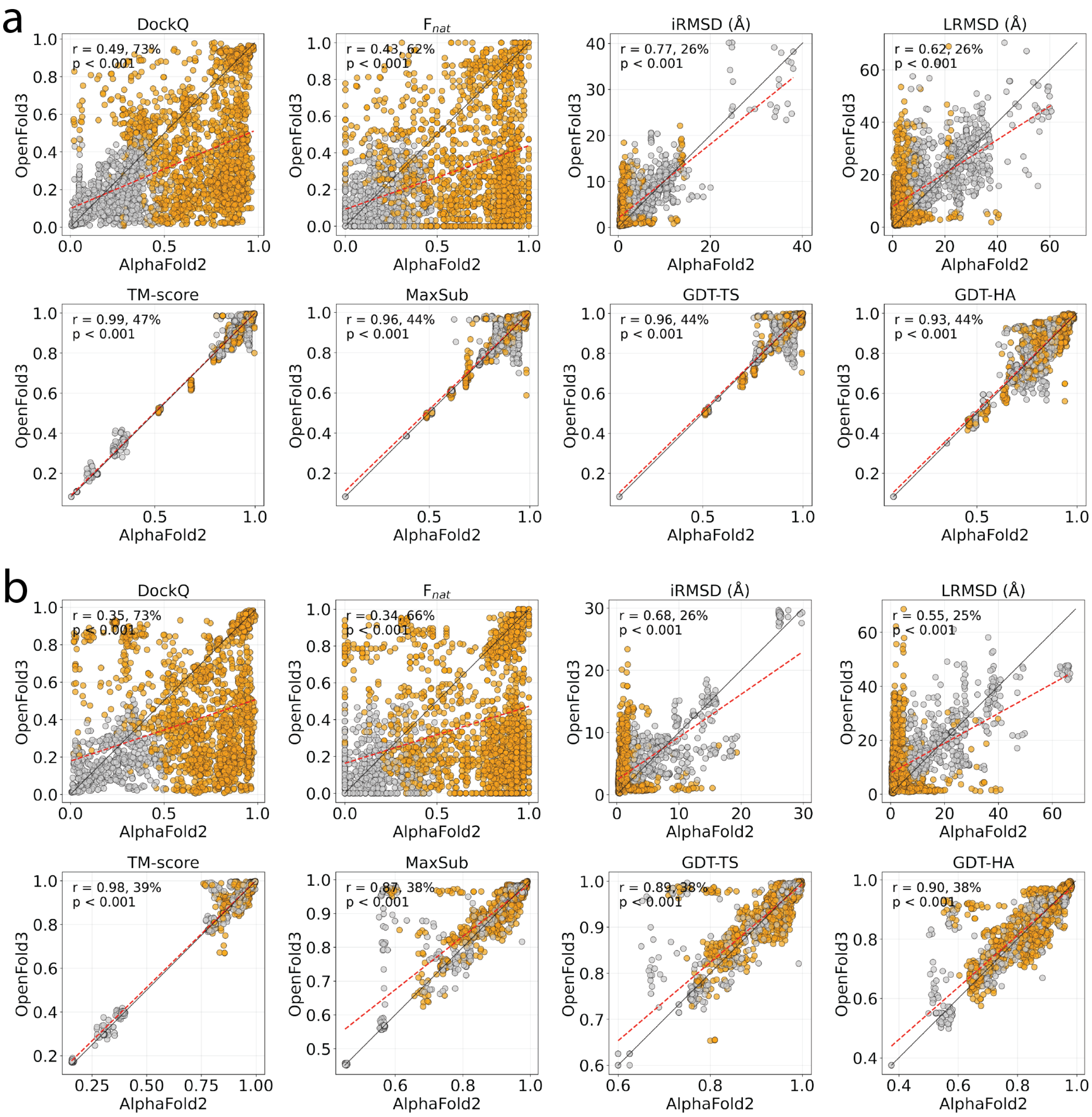


**Fig S3.** AF2 versus OF3 method agreement of structural quality metrics is shown for (a) IDR and (b) Non-IDR subsets. For both panels, top rows show interface metrics of DockQ, $f_{nat}$, lRMSD, and iRMSD, while the bottom rows show global structural similarity metrics. Pearson correlations (r) and proportion of data points with higher AF2 scores than OF3 scores is also given. Gray color indicates incorrect and orange indicates acceptable+ based on CAPRI peptide system.

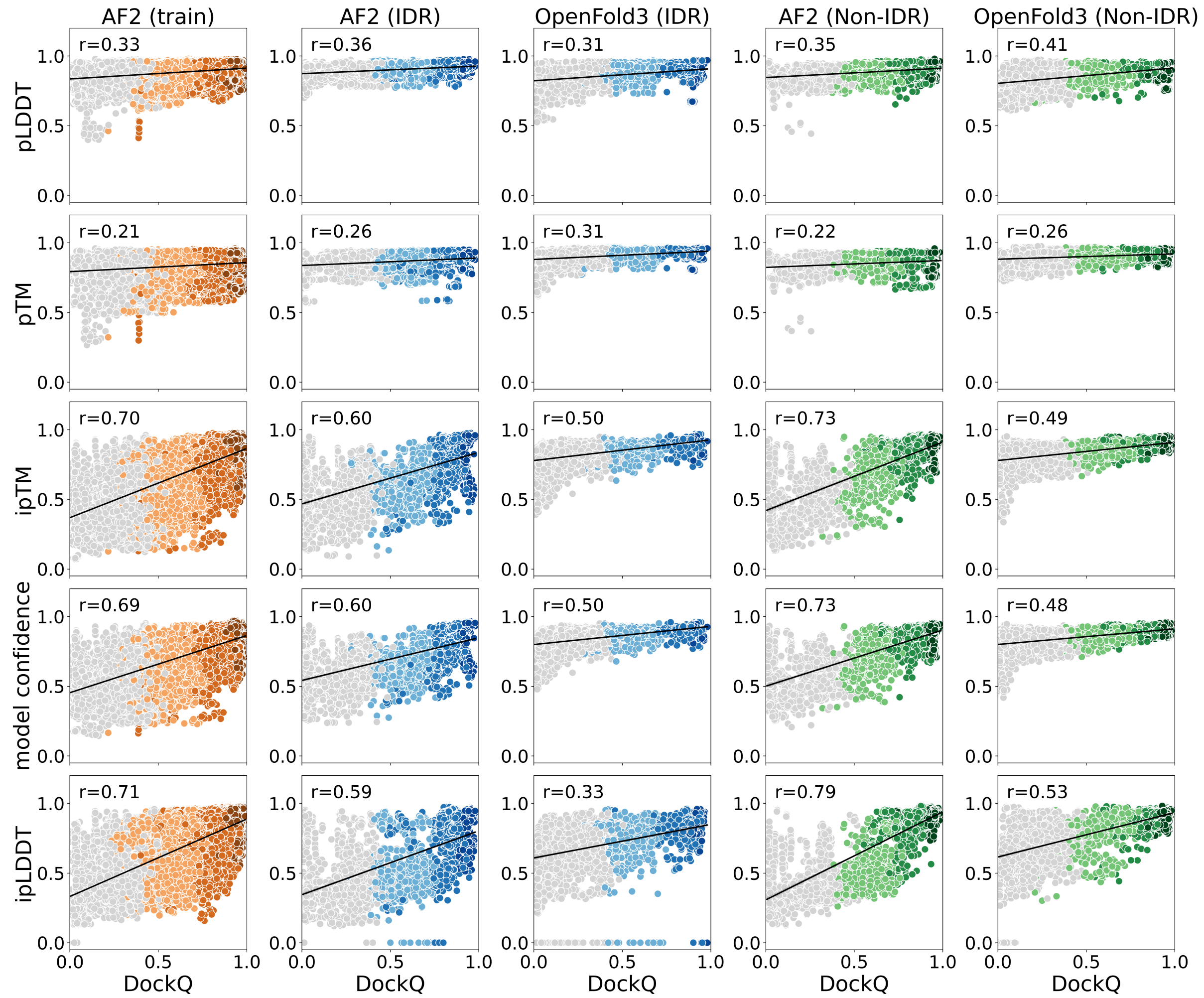


**Fig S4.** Scatter plots of model confidence scores and DockQ. Points are colored according to CAPRI peptide quality categories: gray indicates incorrect, while the gradient from light to dark blue represents acceptable, medium, and high quality classes, respectively. Pearson correlations (r) are reported, all correlations are significant ($p < 0.001$).

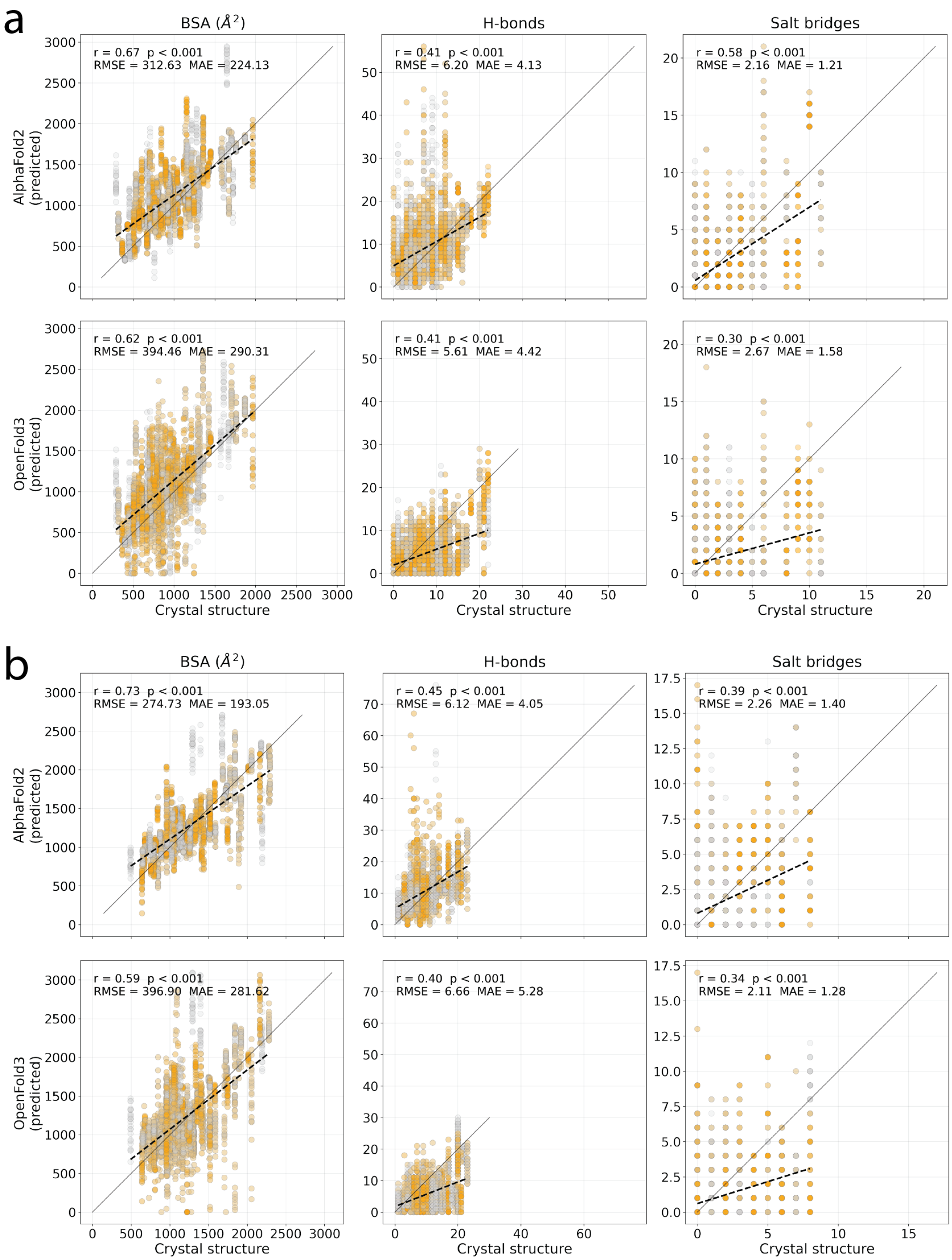


**Fig S5.** Scatter plots comparing predicted versus crystal structure values for three physical properties are shown for (a) the IDR subset and (b) the Non-IDR subset. Each point represents a single prediction, colored gray for incorrect predictions and orange for all other categories based on CAPRI peptide criteria. Black dashed lines denote the least-squares fit; Pearson correlation coefficients (r) with corresponding p-values, RMSE, and MAE are displayed in the upper-left.

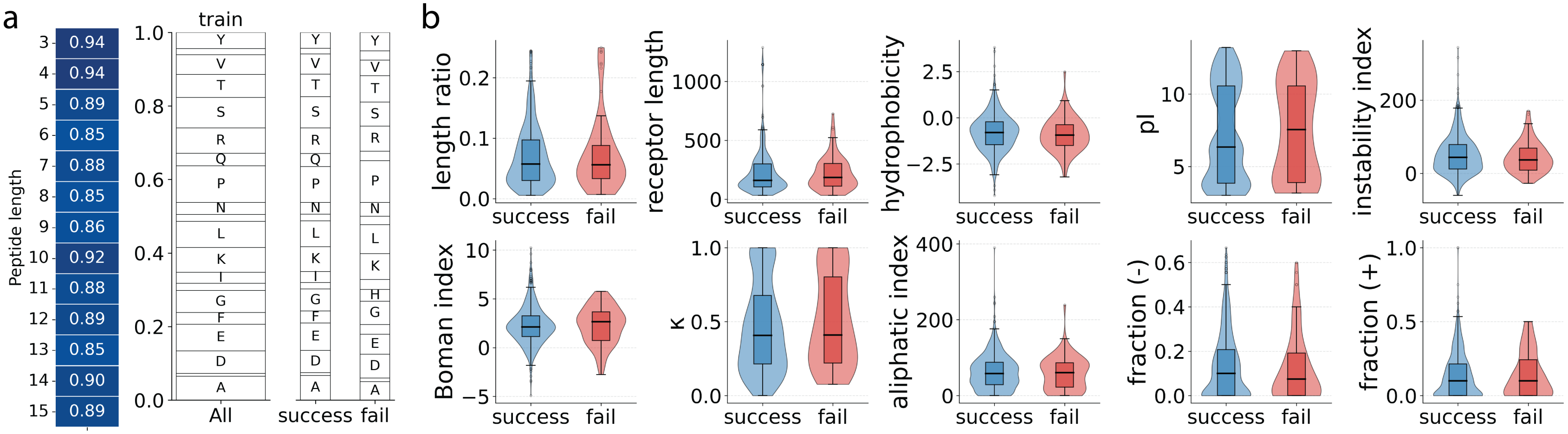


**Fig S6.** (a) Heatmap illustrates the success rates for each peptide length in the training dataset. Amino acid compositions are depicted in stacked bar plots, showing mean fractions in the successful and failed predictions of the training dataset. (b) Distribution of peptide physicochemical features according to prediction outcome. A target is labeled as "successful" if at least one of its 25 predicted models meets acceptable or higher CAPRI peptide criteria; otherwise, it is labeled as "failed". * denote features showing statistically significant differences between successful and failed predictions (Mann-Whitney $U$ test, $p < 0.05$).

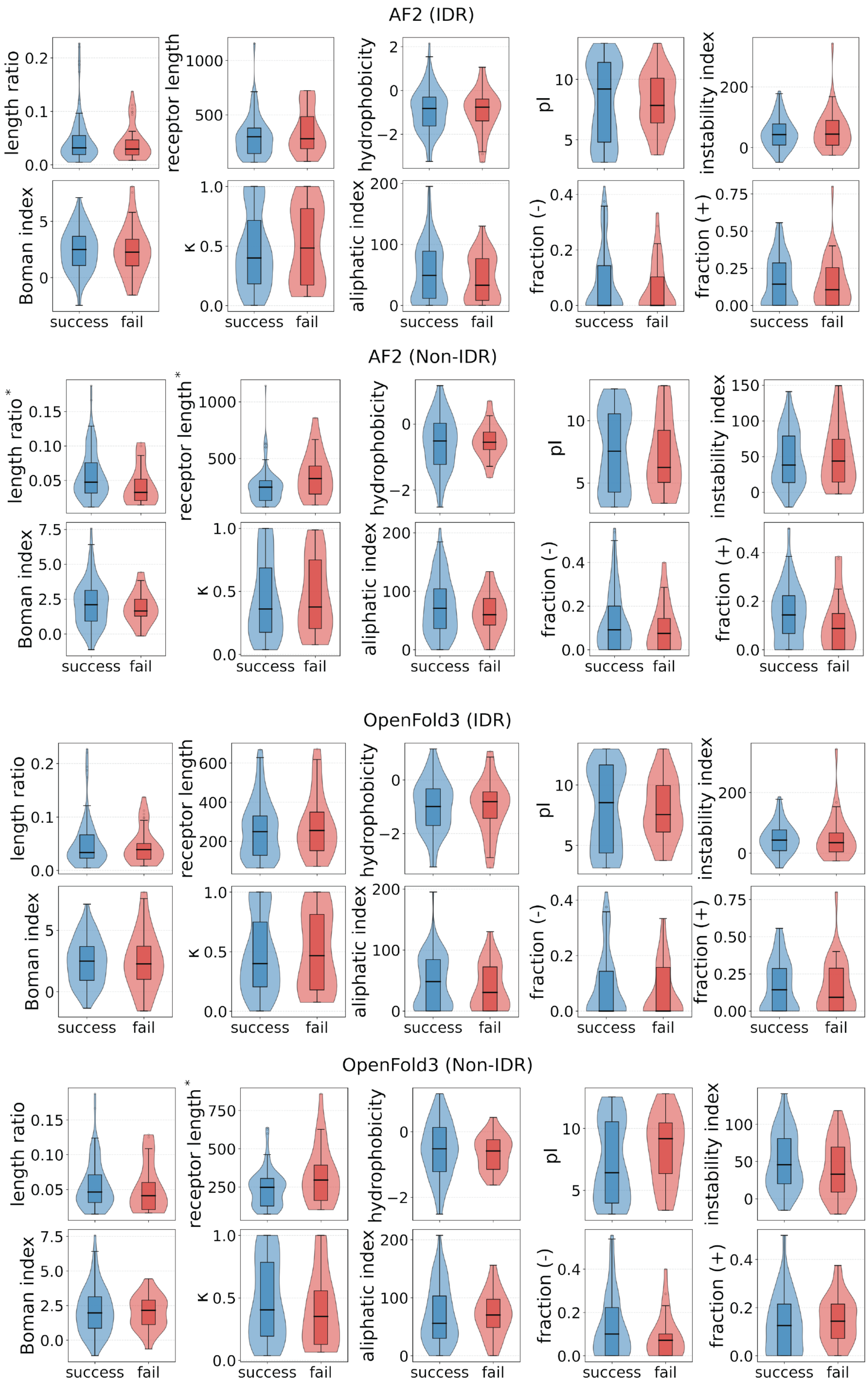


**Fig S7.** Distribution of peptide physicochemical features according to prediction outcome. * denote features showing statistically significant differences between successful and failed predictions (Mann-Whitney $U$ test, $p < 0.05$).
